# Basalin: an evolutionary unconstrained protein revealed via a conserved role in basal plate function

**DOI:** 10.1101/398230

**Authors:** Samuel Dean, Flavia Moreira-Leite, Keith Gull

## Abstract

Most motile flagella have an axoneme that contains nine outer microtubule doublets and a central pair (CP) of microtubules. The CP is thought to coordinate the flagellar beat and defects in CP projections are associated with loss of motility and human disease. In most cilia, the CP nucleate near a ‘basal plate’ at the distal end of the transition zone (TZ). Here, we show that the trypanosome TZ protein ‘basalin’ is essential for building the basal plate, and its loss is associated with inefficient recruitment of CP assembly factors to the TZ, loss of the CP and flagellum paralysis. Guided by synteny, we identified highly divergent basalin orthologs in the genomes of related *Leishmania* species. Basalins are predicted to be highly unstructured, suggesting that they may act as ‘hubs’ facilitating many protein-protein interactions. This raises the general concept that proteins involved in cytoskeletal functions and apparently appearing organism-specific, may have highly divergent and cryptic orthologs in other species.

## Introduction

Flagellar motility is important for cell propulsion and the generation of fluid flow over tissues. The axonemes of most motile flagella have a central pair (CP) apparatus that is comprised of two microtubules, C1 and C2, and their projections (Mitchell & Sale, 1999). The CP apparatus is essential to coordinate the action of dyneins so that the flagellum beat can be propagated efficiently and defects in the CP apparatus cause cell motility defects. For example, loss of Hydin, a component of the C2 projection, causes ciliary bending and beating impairment in mice (Lechtreck, Delmotte, Robinson, Sanderson, & Witman, 2008), defective beat switching in *Chlamydomonas* (Lechtreck & Witman, 2007), and loss of locomotion in trypanosomes (Dawe, Shaw, Farr, & Gull, 2007). Mutations in murine Hydin are associated with hydrocephalus (Davy & Robinson, 2003) and human hydin is also associated with hydrocephalus (Callen, Baker, & Lane, 1990) and primary ciliary dyskinesia (Olbrich et al., 2012).

In contrast to the outer axonemal microtubules, which are extensions of the basal body microtubules, the CP does not elongate directly from the basal body. Rather, the proximal end of one or both CP microtubules is embedded in an electron dense structure at the base of the axoneme termed the basal plate (Höög et al., 2014; May-Simera & Kelley, 2012).

Although the basal plate is ideally located to nucleate the CP, the lack of basal plate mutants or known basal plate components has hampered efforts to determine its function and our knowledge of the role of the basal plate in CP assembly is limited. Ablating trypanosome gamma tubulin results in immotile flagella with no CP (Mckean, Baines, Vaughan, & Gull, 2003), suggesting that CP nucleation is mediated by the gamma tubulin ring complex. In agreement with this notion, gamma tubulin was localised to the flagellum base in trypanosomes (Figure 5, (Scott, Sherwin, & Gull, 1997; Zhou & Li, 2015) and the TZ stellate structure in *Chlamydomonas* (Silflow, Liu, LaVoie, Richardson, & Palevitz, 1999). Knockouts of the microtubule severing protein katanin in *Chlamydomonas* lack the CP (Dymek & Smith, 2012; Dymek, Lefebvre, & Smith, 2004), suggesting a role for microtubule severing in CP assembly. Mating katanin knockout gametes with wild type gametes caused cytoplasmic complementation and suggested CP assembly at sub-distal regions of preformed flagellar axonemes in the resulting zygote, suggesting that CP formation does not require specific nucleating factors to be located at the base of the axoneme (Lechtreck, Gould, & Witman, 2013).

Previously, we reported the development and functional analysis of a trypanosome TZ proteome (Dean, Moreira-Leite, Varga, & Gull, 2016). Here we have analysed the function of a particular TZ protein, that we have named basalin. Mutational analysis shows that RNAi ablation of basalin leads to paralysis and missing CP microtubules. Importantly, the TZs of cells induced for basalin RNAi possess a flagellum but are missing the basal plate, providing insights into the relationship between the basal plate and CP formation. At first sight, and intriguingly for a protein involved in such a conserved function, basalin appeared by homology searches to be a *Trypanosoma brucei* clade-specific protein. However, using syntenic comparison of the genomes we discovered a putative syntenic version of basalin in the *L. mexicana* genome. Deletion of this gene in *L. mexicana* reproduced the missing basal plate phenotype observed upon basalin RNAi in *T. brucei,* suggesting that these proteins are related but extraordinarily divergent in sequence. This establishes that proteins with intrinsic disorder are important for evolutionary conserved cytoskeletal functions and raises the general concept that such proteins involved in cytoskeletal functions and apparently appearing to be organism-specific, may have highly divergent and cryptic orthologs in other species responsible for conserved functions.

## Results

### Basalin knockdown generates flagellum paralysis, short flagella and a cytokinesis defect

Basalin (Tb927.7.3130) was identified as a putative basal plate constituent since it localised to the extreme distal TZ (Dean et al., 2016). We found that when basalin was ablated, the flagella of affected cells were paralysed (supplementary movie 1) and approximately 9 microns shorter than that of uninduced cells (supplementary Figure 1). Cells exhibiting a supernumerary complement of DNA containing structures - the nucleus and the kinetoplast (the mass of mitochondrial DNA) - were also observed (supplementary Figure 1) indicative of a cytokinesis defect and concomitant reduction in population growth. There were a number of associated morphometric effects, such as a pronounced reduction in the distance between the kinetoplast and nucleus, and an increase in the length of the ‘free’ flagellum that extends beyond the anterior of the cell body (supplementary Figure 1). These defects are likely to be consequences of cell division defects caused by the shortened flagellum at cytokinesis.

### Basalin is essential for building the axonemal central pair

To determine the reason for flagellum paralysis after basalin RNAi, we repeated these RNAi induction experiments in cells expressing axonemal marker proteins tagged with a fluorescent protein. We analysed the RNAi phenotype 48 hours after RNAi induction, as this represents the first time-point after induction when basalin was undetectable at the TZ in a large proportion of induced cells, and correlates with the initial observations of morphological aberrations and growth inhibition (supplementary Figure 1). Two components of the CP complex (PF16 and Hydin), a component of the radial spokes (RSP2) and a component of the dynein arms were chosen as markers for this analysis. PF16 (a C1 elaboration) was absent in 63% of cells with a single flagellum after 48 hours RNAi (Figure 1). Dividing cells with an old flagellum that had a strong PF16 signal and a new flagellum that was negative for PF16 were often observed after RNAi, consistent with the pattern of basalin absence from the new flagellum only after RNAi (supplementary Figure 1). Although tagged Hydin (a C2 elaboration) was still detected after basalin RNAi, its signal was fainter and more punctate in 31% of induced cells (compared with the uninduced) (Figure 1). Hydin remained associated with the flagellum even after detergent extraction using a non-ionic detergent, demonstrating that the residual Hydin was stably associated with the flagellum of basalin ablated cells. Flagellum localisation of RSP2 and the axonemal dynein was unaffected by basalin RNAi, suggesting that the radial spokes and dynein arms (doublet microtubules) were not significantly altered.

**Figure 1.**
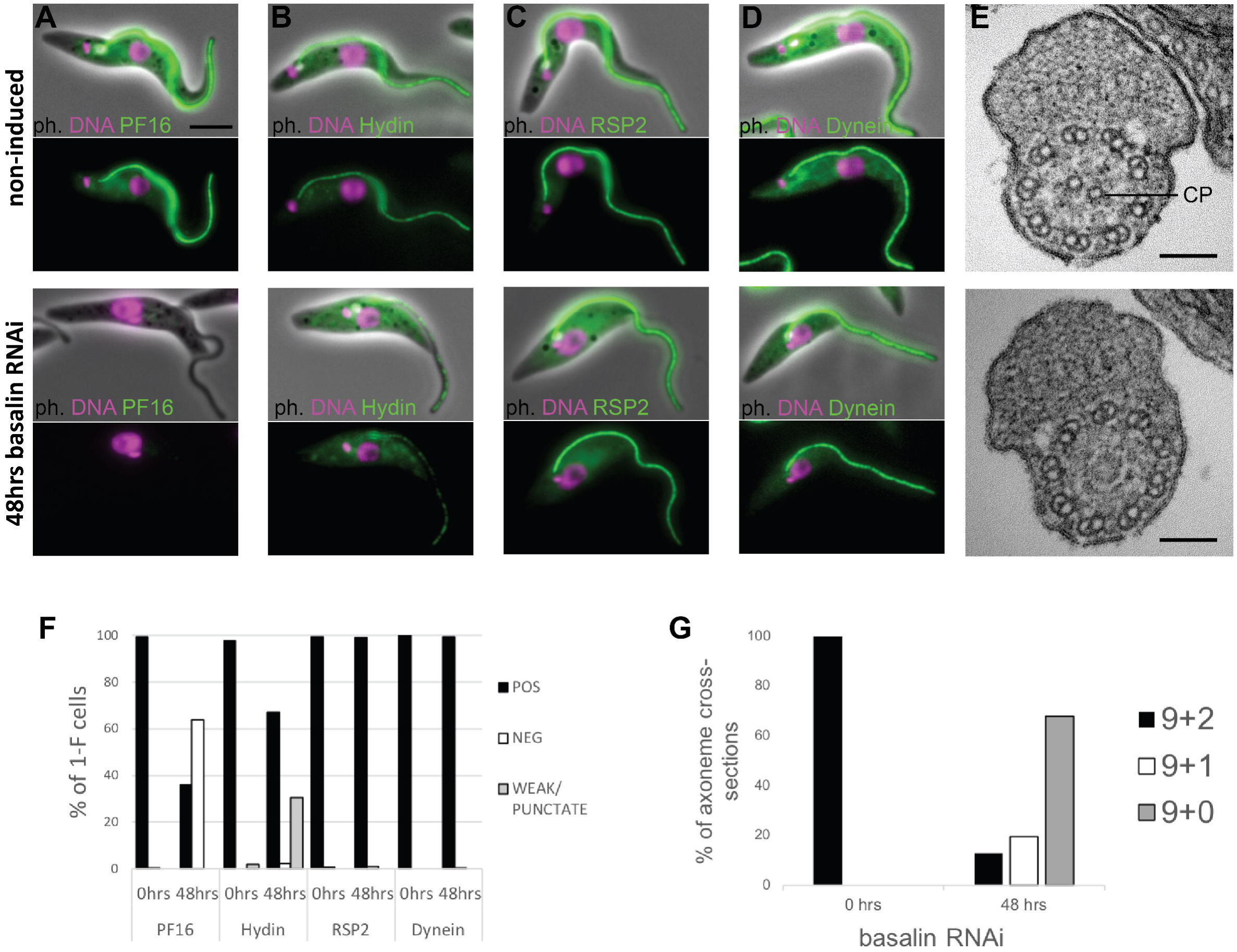
Basalin is essential for building the axonemal central pair. Basalin RNAi was performed in cells expressing tagged markers for different axonemal components, showed that: A) PF16 (Cl projection) is absent from affected flagella, B) hydin (C2 projection) is reduced and punctate, but C) RSP2 (radial spokes marker) and D) dynein (dynein arm marker) flagellar localisation are unaffected. E) TEM cross-sections through the axoneme demonstrate that the central pair microtubules are missing from the axonemes in cells induced for basalin RNAi. F) The presence of axoneme marker proteins (N>200) and G) the number of axonemal microtubules (N=31) were quantified after basalin RNAi.

The flagellum paralysis and PF16 absence suggested a CP defect and hence we analysed the axonemal ultrastructure by transmission electron microscopy (TEM) of cells induced for basalin RNAi. In uninduced cells, all of the axonemal microtubules were clearly visible in all cross-sections of the axoneme, whereas in cells undergoing RNAi for 48 hours, both CP microtubules were missing in 68% of axoneme cross-sections, and 1 CP microtubule was missing in a further 20% of axoneme cross-sections (Figure 1A).

### Basalin is essential for building the TZ basal plate

To study further the effect of basalin RNAi upon TZ structures, we analysed longitudinal sections of the TZ by TEM. In non-induced cells, the characteristic electron density of the basal plate was clearly observed in all longitudinal sections through the centre of the proximal region of the flagellum (comprising the BB, the TZ and the proximal axoneme). However, after 48 hours of basalin RNAi, most longitudinal sections of the proximal flagellum either lacked a visible basal plate (57%) or had a greatly reduced basal plate only (33%) (Figure 2). TZ structures such as the Y linkers and ciliary necklace remained. The proportion of flagella lacking a basal plate correlates with the proportion of axoneme cross-sections with no CP (Figure 1). From this, we conclude that ablating basalin prevents the basal plate from being built and that this is associated with the absence of the CP in most axoneme cross-sections.

**Figure 2.**
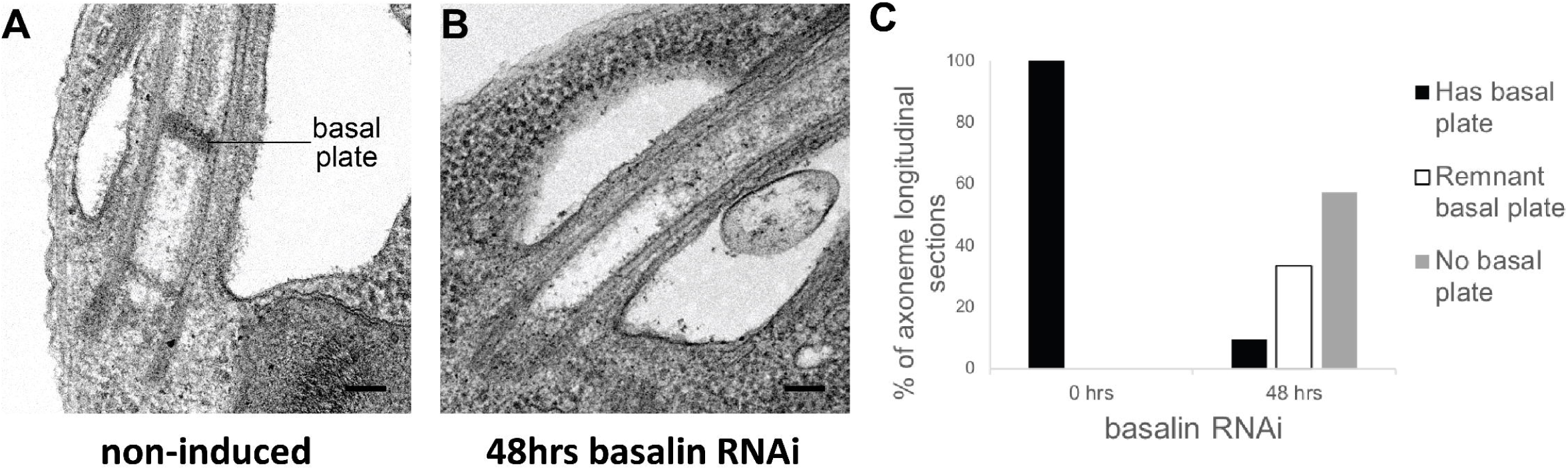
Basalin RNAi produces cells without a basal plate in the flagellum. A) TEM longitudinal sections through the centre of the proximal flagellum (including the transition zone and the proximal end of the axoneme), before (A) and after (B) RNAi. C) The presence and absence of a basal plate in transition zone longitudinal sections was quantified.

### Basalin localises to the basal plate area of the TZ

Basalin was previously localised to the TZ region by eYFP tagging (Dean et al., 2016). We tried immunogold labelling of detergent treated cells (cytoskeletons) mounted onto TEM grids; however, we were not able to obtain reproducible gold labelling, possibly because of embedment of basalin within the basal plate. Hence, to obtain more precise localisation data on basalin, we expressed basalin fused to the bright green fluorophore “mNeonGreen” (Shaner et al., 2013) in a cell line expressing mScarletl∷TZP103.8 (Bindels et al., 2017) and analysed these cells by two-colour microscopy. TZP103.8 (Tb927.11.5770) was for chosen for co-localization because it was previously mapped by two-colour microscopy to a position immediately distal to the basal plate, and it is also required for CP assembly (Dean et al., 2016). Using this method, basalin was localised to a position ˜100nm more proximal than TZP103.8, which corresponds to the expected position of the basal plate. A faint basalin signal was also observed in the pro-basal body (Figure 3A), and this signal increased in intensity as the pro-basal body matured (Figure 3B), eventually becoming visible on both basal body/pro-basal body pairs in late dividing cells (Figure 3C-E). The position of basalin in the expected location of the basal plate, combined with the basal plate absence mutant phenotype, indicate that basalin is an essential component required for basal plate formation and function.

**Figure 3.**
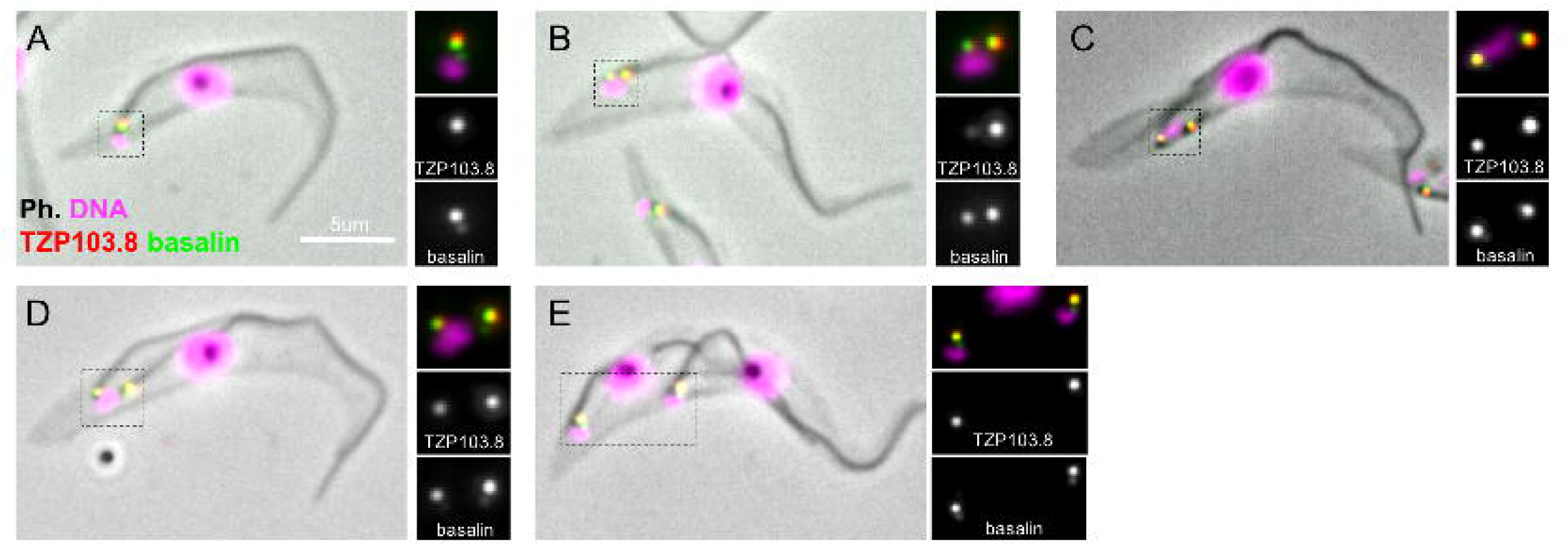
Basalin and TZP103.8 localisation through the cell cycle. A) In non-dividing cells, basalin localises just proximal to the TZP103.8 TZ signal and faintly to the pro-basal body. B) As the pro-basal body matures and moves to the posterior relative to the old flagellum, the basalin signal at the new transition zone increases in intensity and a faint TZP103.8 signal is detected. C) - E) The TZP103.8 signal intensity increases at later stages of the cell cycle, and basalin is detected on the newly formed pro-basal bodies of both old and new flagella.

### Basalin is essential for the efficient recruitment of TZP103.8 to the TZ

Since basalin represents the best candidate for a central component of the basal plate, we therefore next asked what affect ablation of basalin had on recruitment of other proteins which may have a function in the basal plate. Given the similarity in their phenotypes and localisations, we addressed the dependency between basalin and TZP103.8 regarding their recruitment to the TZ. For this purpose, we targeted each individual protein for RNAi separately in a cell line co-expressing fluorescently-tagged copies of both proteins. RNAi targeting basalin was effective as judged by its absence from affected TZs. Strikingly, this resulted in loss of TZP103.8 from affected TZs such that it was either undetectable or almost undetectable by fluorescence microscopy (Figure 4). In contrast, basalin was still recruited efficiently to the flagellum when TZP103.8 was ablated (Figure 4). Importantly, TZP103.8 RNAi, whilst resulting in a lack of CP microtubules did not cause the disappearance of the basal plate (Dean et al., 2016). Basalin and TZP103.8 remained associated with isolated flagella, even after extraction with near-saturating concentrations of sodium chloride or potassium chloride (data not shown), suggesting they are core structural components of these structures.

**Figure 4.**
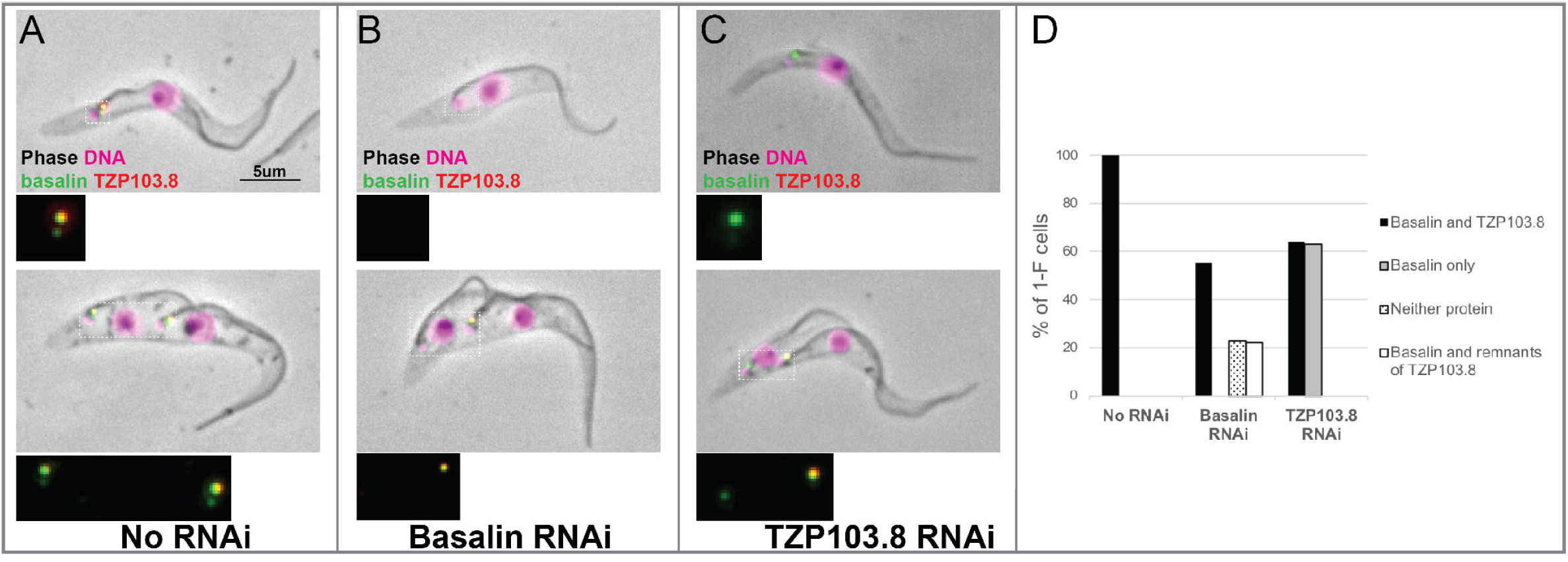
TZP103.8 depends on basalin for its localisation to the transition zone. Basalin or TZP103.8 were ablated by RNAi in cells co-expressing mScarlet∷TZP103.8 (red) and mNeonGreen∷basalin (green) to determine their dependency relationship at the transition zone. A) In uninduced cells, basalin and TZP103.8 were both detected at the transition zone. B) When basalin was ablated by RNAi, TZP103.8 was not efficiently recruited to the transition zone. C) In contrast, when TZP103.8 was ablated, basalin still localised to the transition zone. Top panels show a cytoskeleton from a non-dividing cell with the flagellum absent for the targeted protein, panels show cytoskeletons from dividing cells with the new flagellum absent for the protein being targeted, bottom. D) Quantification of basalin/ TZP103.8 presence at the transition zone in non-induced cells and after 48 hours RNAi of either protein confirms the differential dependencies at the transition zone.

**Figure 5.**
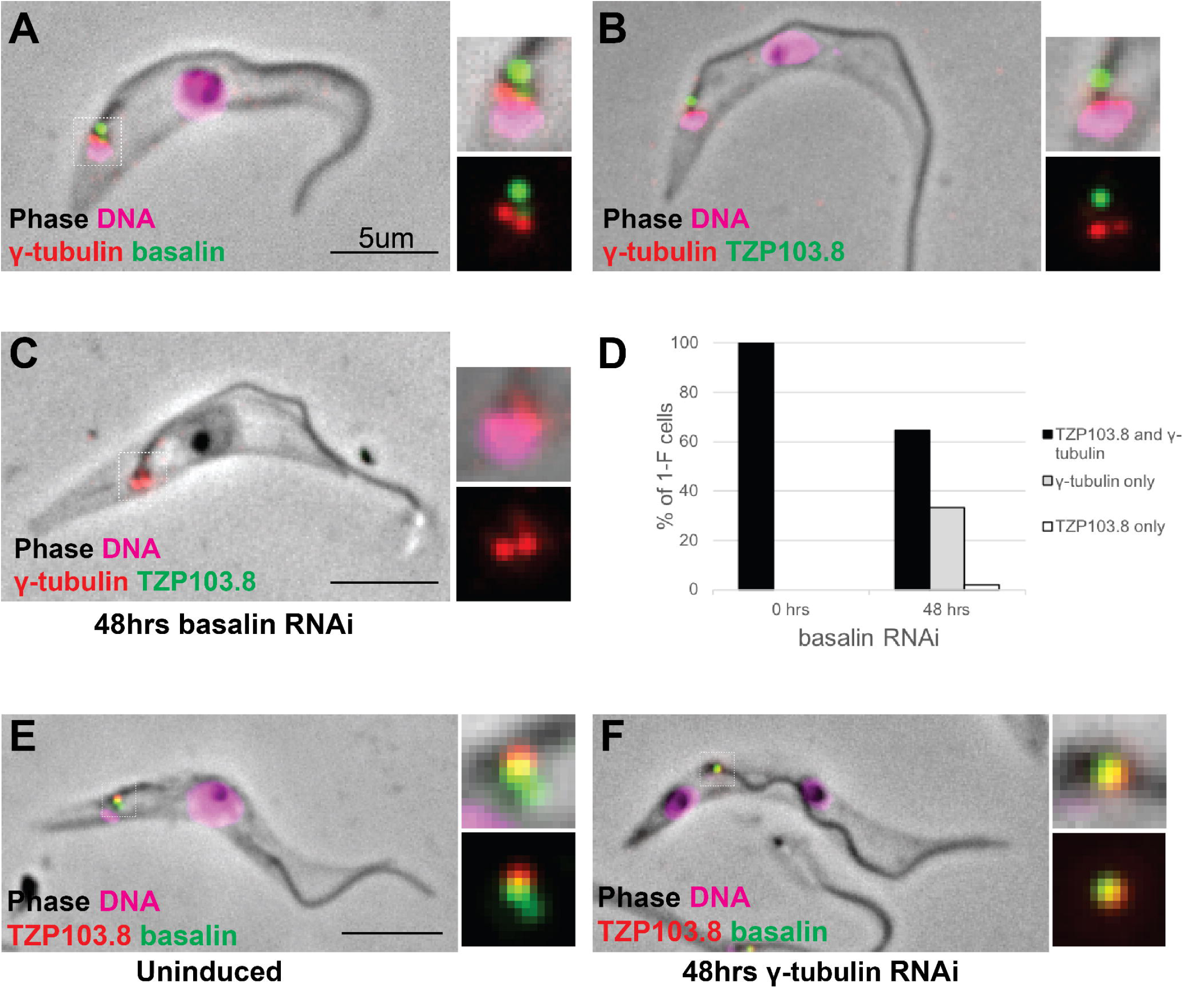
Co-localisation and dependency of gamma tubulin with basalin and TZP103.8. Gamma tubulin localises to the base of the flagellum more proximal to A) basalin and B) TZP103.8. C) and D) Ablating basalin does not prevent gamma tubulin from being recruited to the base of the flagellum. E) and F) Ablating gamma tubulin does not prevent basalin or TZP103.8 from being recruited to the TZ.

### Gamma tubulin basal body localisation is independent of basalin and TZP103.8 TZ localisation

Gamma tubulin has an essential role in assembling the axonemal CP (Mckean et al., 2003; Zhou & Li, 2015). Therefore, we investigated whether there was a functional link between gamma-tubulin localization and the recruitment of basalin or TZP103.8 to the TZ. Firstly, we expressed gamma tubulin tagged with mNeonGreen in cells expressing either basalin or TZP103.8 tagged with mScarlet (Figure 5). Gamma tubulin localised ˜400nm proximal to basalin and TZP103.8, with no overlap of signal between gamma tubulin and these TZPs. This is consistent with previous data that localised components of the gamma tubulin ring complex to the basal body area in trypanosomes (Zhou & Li, 2015). RNAi targeting basalin caused a general reduction in the gamma tubulin signal, but did not inhibit the recruitment of gamma tubulin to the basal body (Figure 5C and D). Similarly, although RNAi targeting gamma tubulin resulted in the expected paralysed flagellum phenotype (data not shown), it did not prevent recruitment of basalin or TZP103.8 to the TZ (Figure 5E and F).

### Basalin has a highly divergent syntenic ortholog in *Leishmania mexicana*

Initial bioinformatics analyses using homology search approaches such as Orthofinder (Emms & Kelly, 2015) suggested that basalin had a highly restricted evolutionary distribution and was not detected outside the *Trypanosoma spp.* These results were unexpected, given the widespread evolutionary presence of a flagellum/cilium basal plate and the fundamental and conserved nature of CP nucleation within motile cilia and flagella. Other kinetoplastid species have a motile flagellum in at least one lifecycle stage and one might therefore expect to detect a conserved protein ortholog. However, sequence analysis even failed to detect an ortholog of basalin in the *Leishmania* species, which are separated from *T. brucei* by only 300 million years (Overath, Haag, Lischke, & O’hUigin, 2001; Stevens & Gibson, 1999; Stevens & Rambaut, 2001). Moreover, even in the very closely related *Trypanosoma congolense* the ortholog appeared very highly divergent to that in *T. brucei* (supplementary Figure 2). A striking property of most of the kinetoplastid genomes available (including that of trypanosomes and *Leishmania)* is that they are highly syntenic, which may be linked to their polycistronic transcription. Genome sequencing has revealed that kinetoplastid genes are arranged in directional gene clusters and that, even when there is extensive sequence divergence, synteny is highly conserved (Ghedin et al., 2004). *Leishmania spp.* are kinetoplastid parasites that can also build a motile flagellum with an axonemal CP and a discrete basal plate in the distal TZ. Thus, we searched for a *L. mexicana* ortholog of basalin by scanning the syntenic genomic region.

Intriguingly, although gene synteny either side of the basalin gene is highly conserved between *T. brucei* and *L. mexicana,* the syntenic locus of basalin in *L. mexicana* is occupied by LmxM.22.1070, which is not a clear sequence homolog of basalin (Figure 6). However, we re-examined the BLASTP scores between *T. brucei* and *L. mexicana* and noticed that LmxM.22.1070 was indeed the reciprocal best BLASTP hit for basalin in *L. mexicana,* but the basalin-LmxM.22.1070 reciprocal BLASTP homologies were extremely weak (e= 0.27 and 1.9, respectively). An HHpred-driven protein comparison (Söding, 2005; Zimmermann et al., 2018) between basalin and LmxM.22.1070 demonstrated that most of the conserved residues lie in, or near, predicted helices (supplementary Figure 2). Although neither protein contained conserved domains that could provide functional clues, some coiled-coils were detected and, importantly, both proteins were predicted to be ˜65% intrinsically disordered (supplementary Figure 2). We therefore set out to determine whether basalin and LmxM.22.1070 were divergent syntenic orthologs. To this end, we expressed tagged LmxM.22.1070 in *L. mexicana,* and demonstrated that this protein localised to the basal body and pro-basal body areas of the flagellum (Figure 7). RNAi is not possible in *L. mexicana* and so we produced LmxM.22.1070 knockouts generated using CRISPR/CAS9 (Beneke et al., 2017). These were viable but had substantial morphological aberrations (supplementary Figure 3). Similar to basalin RNAi cells, LmxM.22.1070 knockout cells had a strong motility defect (supplementary movie 2) slower doubling times and substantially shorter flagella, with most cells having no ‘free’ flagellum extending from the flagellar pocket (the cell surface invagination from which the flagellum emerges). Knockout cells were shorter and more round than the parental cells, consistent with the important link between flagellum biogenesis and cell shape (Sunter & Gull, 2017).

**Figure 6.**
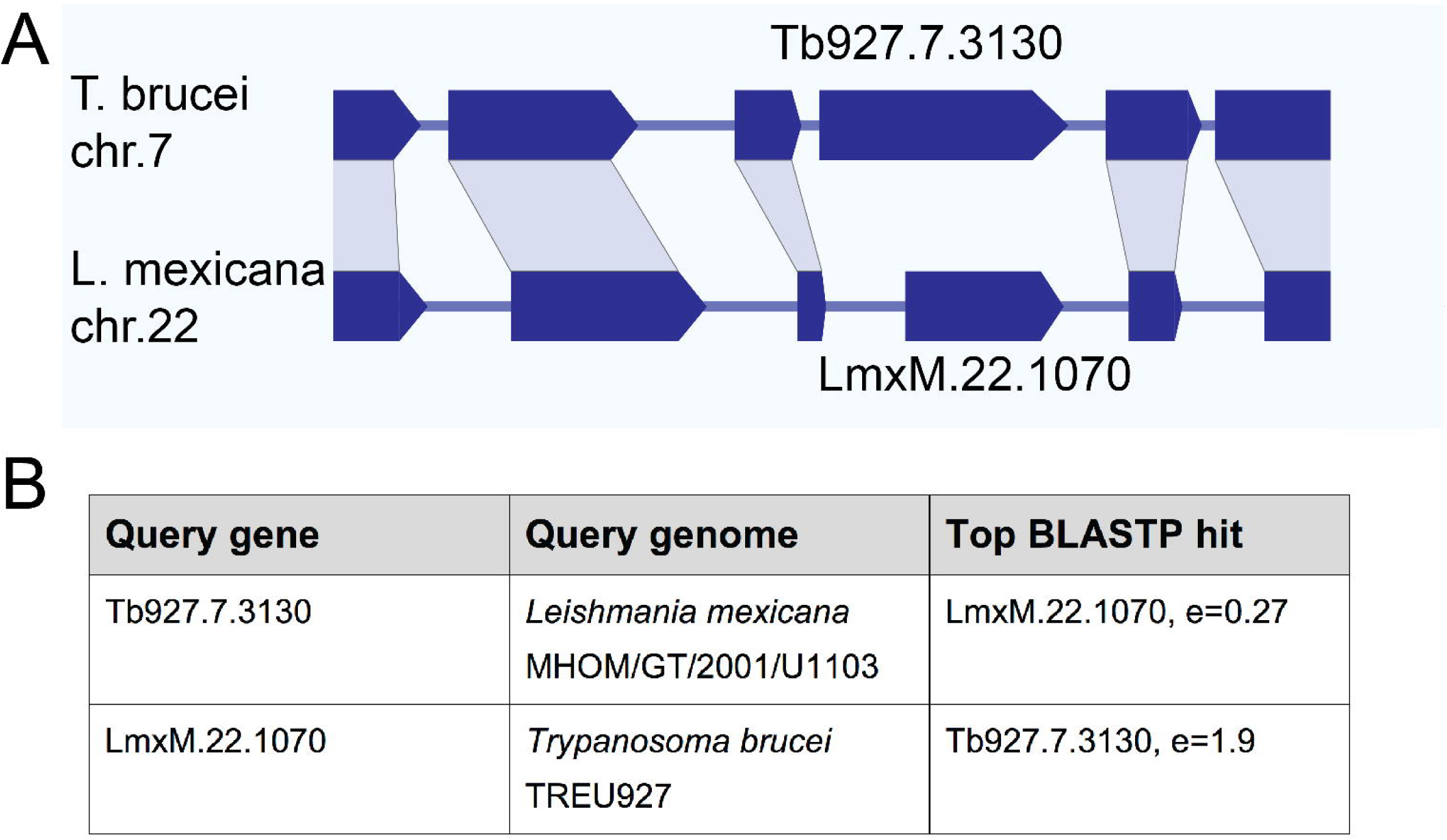
A) Basalin and LmxM.22.1070 genes are at syntenic positions along chromosome *Leishmania mexicana* chromosome 22 and *Trypanosoma brucei* chromosome 7, respectively. B) Basalin and LmxM.22.1070 are very weak reciprocal best BLAST hits.

**Figure 7.**
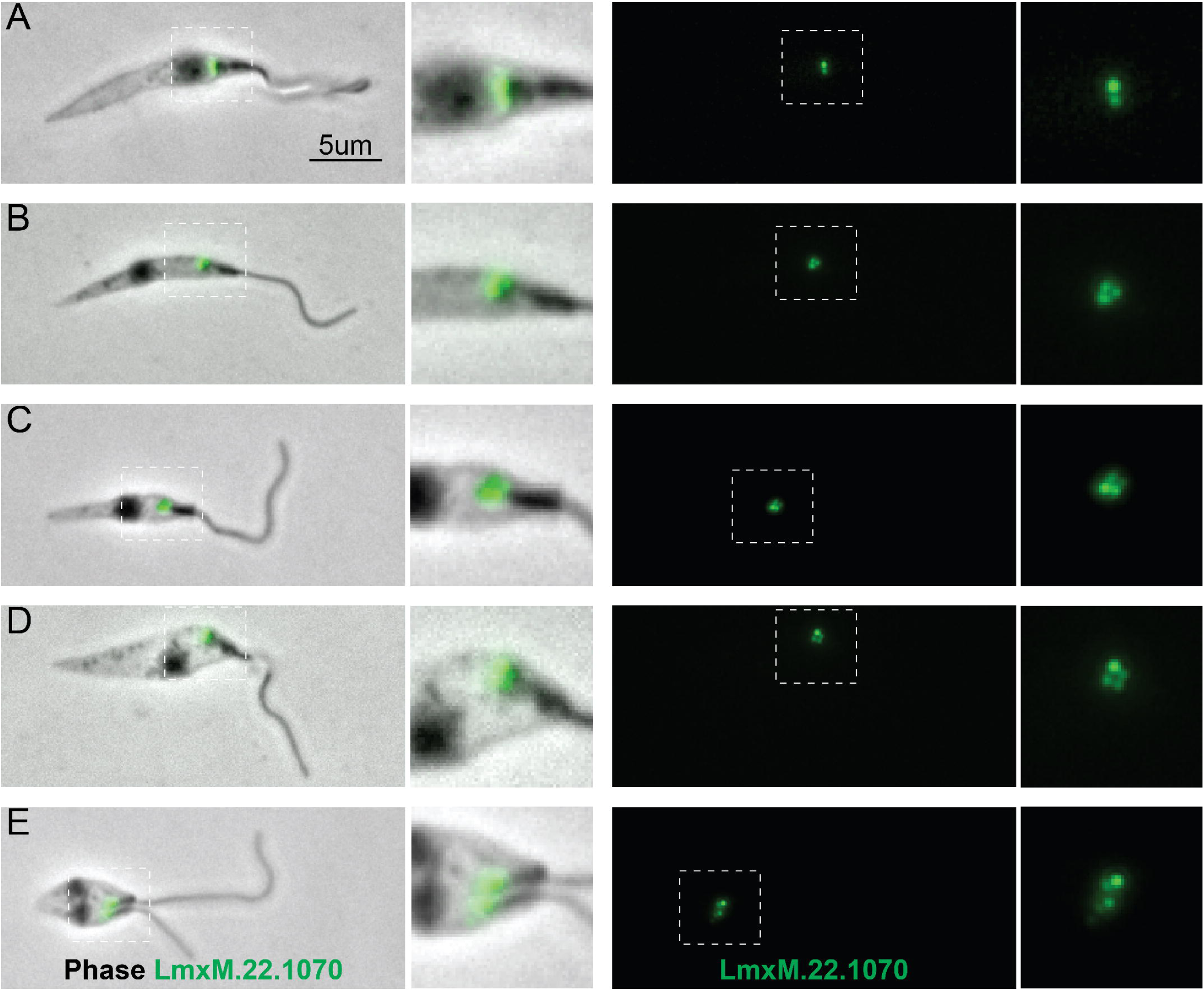
Localisation of LmxM.22.1070 in *L. mexicana* promastigotes through the cell cycle recapitulates that of basalin in trypanosomes. A) In non-dividing cells, LmxM.22.1070 localises to a position consistent with the transition zone and the pro-basal body. B) - E) As the cell divides, the basal bodies are duplicated and LmxM.22.1070 is detected on the basal body and pro-basal body of both flagella in dividing cells.

EM analysis demonstrated that, as with basalin RNAi in trypanosomes, the CP was absent from a large proportion (41%) of the axonemal cross sections examined (Figure 8). Importantly, we could not detect a basal plate or an associated CP in any longitudinal sections through the TZ in LmxM.22.170 knockout cells or in an EM tomogram (Supplementary Figure 3), while a basal plate was nearly always observed at the TZ of parental cells (Figure 8). The morphological and growth defects were largely rescued by LmxM.22.1070 episome-based expression, but not by *T. brucei* basalin expression using the same system (supplementary Figure 3).

**Figure 8.**
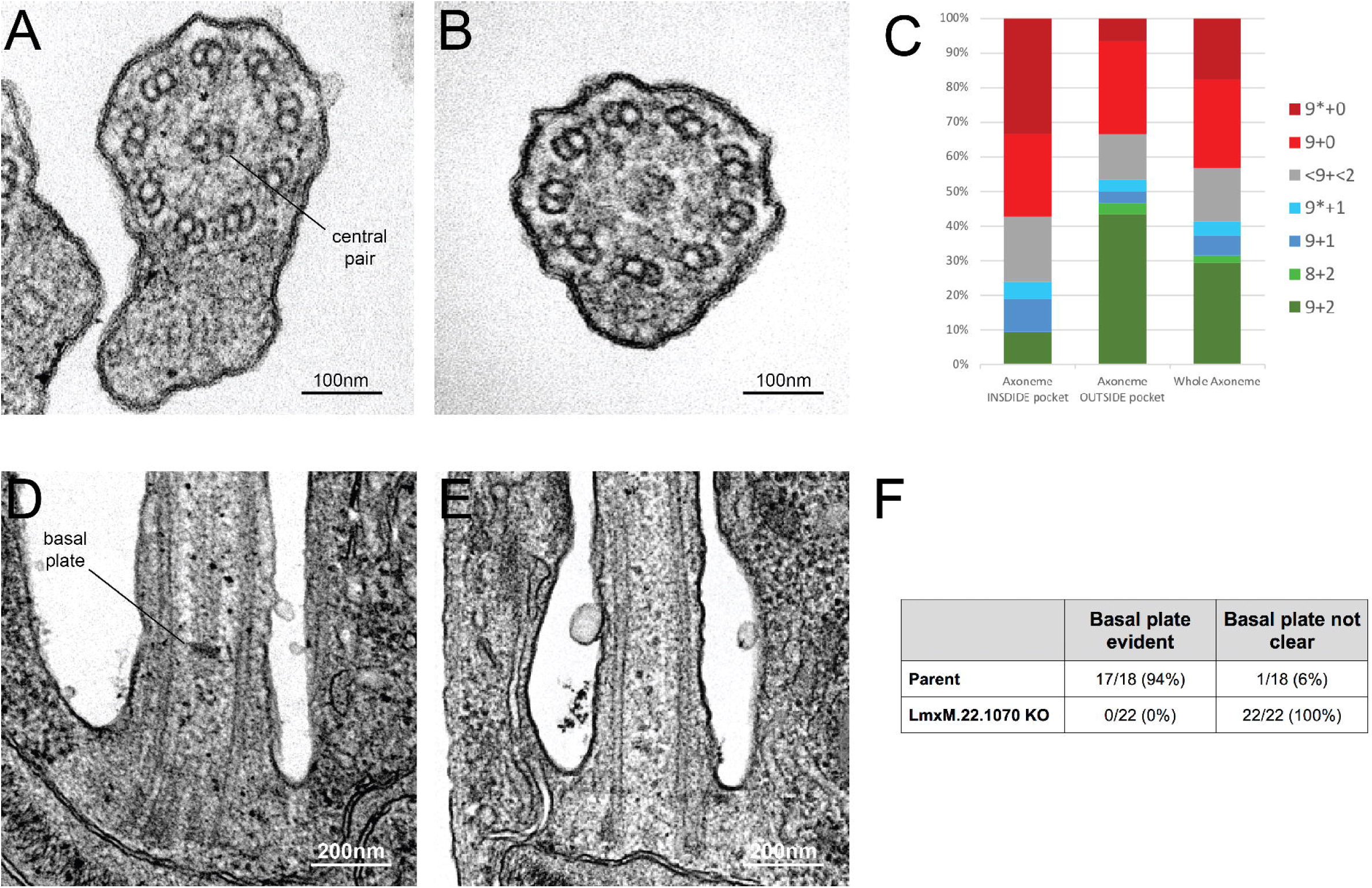
Knockout of LmxM.22.1070 in *L. mexicana* promastigotes causes flagella to be built without the central pair microtubules and a basal plate. A) and B) TEM cross sections through the axoneme show that the central pair is absent in knockout cells. C) Quantification of the microtubules arrangement in KO axonemes shows that >40% of cross sections are missing their CP, and that this is phenotype is more penetrant in the proximal axoneme. The asterisk refers to the microtubules being disorganised. D) and E) TEM longitudinal sections through the transition zone demonstrate that the basal plate is absent. F) Quantification shows that the basal plate was not observed any the transition zones that were examined in KO cells.

Therefore, a combination of bioinformatics, localisation and functional evidence demonstrates that LmxM.22.1070 is a highly divergent syntenic ortholog of basalin. Basalin is “conserved” in the kinetoplastids and possibly in many other organisms where it remains cryptic because of extreme sequence diversity and lack of clues from synteny.

## Discussion

In this work, we describe basalin (Tb927.7.3130), a protein identified in our trypanosome flagellum transition zone proteome (Dean et al., 2016), and show that it is required for construction of the basal plate of the flagellum. Whilst a shorter axoneme is built in cells in which basalin was ablated, the absence of the basal plate leads a strong defect in CP assembly and flagellum immotility. Basalin appears to be fundamental for basal plate formation and to our knowledge this is the first example of a protein required for basal plate formation. Moreover, mutant analysis clearly demonstrates the link between the TZ basal plate structure and CP assembly. Basalin is required for recruitment of another, unrelated protein, TZP103.8, also required for CP formation. However, gamma tubulin, which again is required for CP formation, can still associate with the basal body in the absence of basalin, suggesting complex pathways are involved in CP formation. It is unlikely that the alternative view that the lack of a CP causes basal plate absence is correct since the basal plate remains in other mutants that lack the CP, such as TZP103.8 (Dean et al., 2016) and PF16 (Beneke et al., 2017; Branche et al., 2006) ablation mutants.

Although all examined TZs of *Leishmania* LmxM.22.1070 KO cells were missing their basal plate and the associated CP, 31% and 10% of axonemal cross-sections revealed a 9+2 and 9+1 axoneme, respectively. How are these VCPs formed in the absence of a basal plate? It is obviously unlikely that the CP that was observed in these cross-sections extends from a basal plate (p<0.02) because no basal plates were observed in the KO TZ longitudinal sections. Similarly, it is unlikely that the CP extends from a plate-less TZ (p<0.02) because no CP was observed extending from the expected position of the basal plate in TZ longitudinal sections. Moreover, cross-sections of distal axonemes (from outside the flagellar pocket) were twice as likely to contain at least one CP microtubule as cross-sections of proximal axonemes (sections inside the flagellar pocket) and an EM tomogram of an entire short axoneme did not reveal any evidence of a CP. All of this evidence, taken together suggests that, when it occurs, CP nucleation happens in the distal axoneme in KO cells. Therefore, the observed CPs may reflect stochastic self-assembly of the CP microtubules inside axonemes, possibly due to mis-localisation of nucleating factors in the absence of the basal plate. Consistent with this, studies in *Chlamydomonas* demonstrate that the CP can self-assemble at different points along pre-formed flagella during cytoplasmic complementation of katanin mutants (Lechtreck et al., 2013). Under the normal conditions of an elongating flagellum, the presence of a basal plate located nucleating factor may facilitate the assembly of a full CP running from the basal plate to tip.

Gamma tubulin and the gamma tubulin ring complex are essential for CP nucleation in trypanosomes (Mckean et al., 2003; Zhou & Li, 2015) but the mechanism by which this occurs is not clear. Our data using fluorescent protein tagging, and previous findings using small epitope tags, localised gamma tubulin to the proximal end of the basal body, at a position ˜400nm proximal to the site of CP nucleation. A clear structural ortholog of the normal cilium TZ basal plate is not evident in the *Chlamydomonas* flagellum, but gamma tubulin has been located to the stellate structure within its TZ (Silflow et al., 1999) which suggests a possible analogous role to the basal plate in CP assembly. The concentration of gamma tubulin signal at the trypanosome basal body may represent a pre-assembly pool, possibly interacting with alpha and beta tubulins and cofactors for more distal CP assembly. Alternatively, there could be a small, undetected focus of gamma tubulin within the TZ basal plate area, although other proteins can be easily located at the basal plate. Thus, gamma tubulin may be nucleating the CP directly at the basal plate, or if the more delocalised concentration of gamma tubulin at the basal body proximal end is real then this may indicate that CP nucleation takes place at a slight distance from the basal plate. In this latter model the basal plate may act to capture a CP structure whose assembly is initiated elsewhere, rather than nucleating the CP. However, in both the nucleating and capturing models there must be number control, i.e. two and only two CP microtubules are incorporated into the centre of the axoneme. This control could be exercised by either the number of nucleation sites or the number of capturing sites within the basal plate.

At the base of the flagellum in *L. mexicana* parental cells, the “lumen” of the transition zone has a different electron density from that of the inner region of the axoneme, with the basal plate delimiting a clear “border” between the TZ and axoneme inner regions. In contrast, longitudinal sections through the centre of the proximal flagellum of basalin KO cells revealed no clear border between the inner regions of the TZ and the axoneme, as these regions had a homogeneous electron density.

Initial analyses failed to find basalin orthologs outside the *Trypanosoma spp.* and it was only after rigorous examination of the synteny, BLAST score, localisation and function that we were able to conclude the LmxM.22.1070 is the *L. mexicana* ortholog of basalin. Interestingly, both TbBasalin and LmexBasalin are predicted to contain ˜65% intrinsically disordered residues. Proteins with a high degree of intrinsic disorder have been shown to associate with different elements of the cytoskeleton, including microtubules, actin, intermediate filaments and motor proteins (reviewed in (Guharoy, Szabo, Contreras Martos, Kosol, & Tompa, 2013)). Their roles include acting as hub proteins in protein-protein interaction networks (Dosztányi, Chen, Dunker, Simon, & Tompa, 2006; Dunker, Cortese, Romero, Iakoucheva, & Uversky, 2005), scaffold proteins that cause “enforced proximity” of proteins to promote their interaction (Buday & Tompa, 2010), or acting as molecular ‘glue’ that becomes rigid once the protein complex has formed (Uversky, 2015). Often only a few specific peptides are conserved and these proteins acquire structure upon interaction with a binding partner (Cumberworth, Lamour, Babu, & Gsponer, 2013; Davey et al., 2012). Therefore, it may be that basalin is under little selective pressure to retain its primary sequence and that a few specific residues in a disordered polypeptide are sufficient to retain its interactions and function. Supporting this, the HHpred-driven alignment of TbBasalin and LmexBasalin suggests that a core helical scaffold is conserved that is rather plastic to insertions of sequences predicted as ‘disordered’. Nonetheless, it is worth noting that the expression of TbBasalin in *L. mexicana* did not rescue the LmexBasalin knockout phenotype, demonstrating some degree of biochemical divergence.

Given the difficulties in identifying the relationship between the trypanosome and *Leishmania* orthologs, more distant orthologs in other organisms would not be detected using BLAST-based methods. This raises the intriguing possibility that basalin has distant orthologs in other organisms and represents a key player in a fundamental mechanism of basal plate formation and CP assembly. The discovery that a fundamental and conserved flagellum structure and function involves such widely divergent orthologues was totally dependent on our use of kinetoplastids such as *Leishmania* and trypanosomes, whose genomes have retained an enormous degree of synteny. This is one of the few groups of organisms where this type of syntenic genome analysis is possible. This raises the fundamental issue that the proportion of lineage-specific proteins in flagellar proteomes of the various model systems is likely to have been over-estimated, and that some proteins that have been categorised as clade specific may, in fact, be highly divergent ancient proteins.

## Materials and methods

### Light and electron microscopy

Live cells and detergent extracted ‘cytoskeletons’ were prepared as described (Dean et al., 2015). Images were acquired using a Leica wide-field fluorescence microscope with a 63x 1.4 NA objective using an sCMOS camera (Andor) and analysed using FIJI. Where the purpose was to determine the relative localisation of proteins within the cell, the microscope’s chromatic aberration was measured using TetraSpec beads (Thermofisher) and then automatically corrected using a custom python script executed in FIJI. DNA was visualized by treating cells or cytoskeletons with 1 μg.mL^−1^ Hoechst 33342. Time lapse movies were captured using an Olympus CKX41 inverted phase-contrast microscope and a Hamamatsu ORCA-ER camera using 100ms exposure at maximum frame rate.

### Cell culture

Trypanosome SMOX 927 cells (Poon, Peacock, Gibson, Gull, & Kelly, 2012) and *L. mexicana* CAS9/T7 cells (Beneke et al., 2017) were cultured in SDM79 for all experiments (Brun & Schönenberger, 1979). Trypanosome and *Leishmania* cell lines were generated as described (Beneke et al., 2017; Dean et al., 2015). RNAi was induced using 2 μg.mL^−1^ doxycycline. Cell densities were measured using a CASYcounter.

### Molecular cloning

Primer annealing sites for making RNAi constructs were designed by RNAit (Redmond, Vadivelu, & Field, 2003) and the resulting amplicon was cloned into the RNAi stemloop vector pQuadra (Inoue et al., 2005). Primers for tagging trypanosome proteins were designed using TAGit (Dean et al., 2015) and primers for generating *Leishmania* knockout cell lines were designed using LeishGEdit (Beneke et al., 2017).

### Bioinformatics

Evolutionary analyses was initially based on Orthofinder output (Emms & Kelly, 2015) with subsequent refinement using BLASTP and gene synteny (Aslett et al., 2010). Various algorithms were used to predict protein alignment (Mergealign (Collingridge & Kelly, 2012), HHpred (Söding, 2005; Zimmermann et al., 2018)), coiled-coils (Pair2 (McDonnell, Jiang, Keating, & Berger, 2006)) and intrinsic disorder (PrDOS (Ishida & Kinoshita, 2007)). Synteny was calculated using TriTrypDB’s (Aslett et al., 2010; Aurrecoechea et al., 2017) inbuilt OrthoMCL with a le^-5^ cut-off and the graphic reproduced using Adobe Illustrator.

### Supplemental material

Figs S1 describes the quantitation of basalin knockdown and phenotype in trypanosome cells. Fig S2 is a bioinformatics analysis comparing basalin with its *L. mexicana* ortholog. Fig S3 is a phenotypic analysis of *Leishmania mexicana* LmxM.22.1070 knockout cells. Supplementary movie 1 shows trypanosome cells undergoing basalin RNAi. Supplementary movie 2 shows LmxM.22.1070 knockout *Leishmania mexicana* cells.

## Acknowledgements

Work in the K.G. laboratory is funded by the Wellcome Trust (WT066839MA and 104627/Z/14/Z). We thank Professor Matthew Higgins for his advice on basalin structure, Dr Fernando Bazan for his advice on basalin alignments, Dr Susanne Warrenfeltz and Dr Omar Harb from EuPathDB for their help on basalin synteny, and Tom Beneke for his advice on *Leishmania* CRISPR. The authors declare no competing financial interests.

## Author contributions

SD: designed the study, performed the experiments, analysed the data, wrote the paper. FM-L: performed the experiments, analysed the data and advised on the manuscript. KG: advised on experiments and writing the manuscript.

---

Supplementary movie 1.

An RNAi screen for paralysed flagella identifies basalin.

Supplementary movie 2.

LmxM.22.1070 knockout *Leishmania mexicana* cells are mostly paralysed with little or no flagellum visible.

**Supplementary Figure 1.**
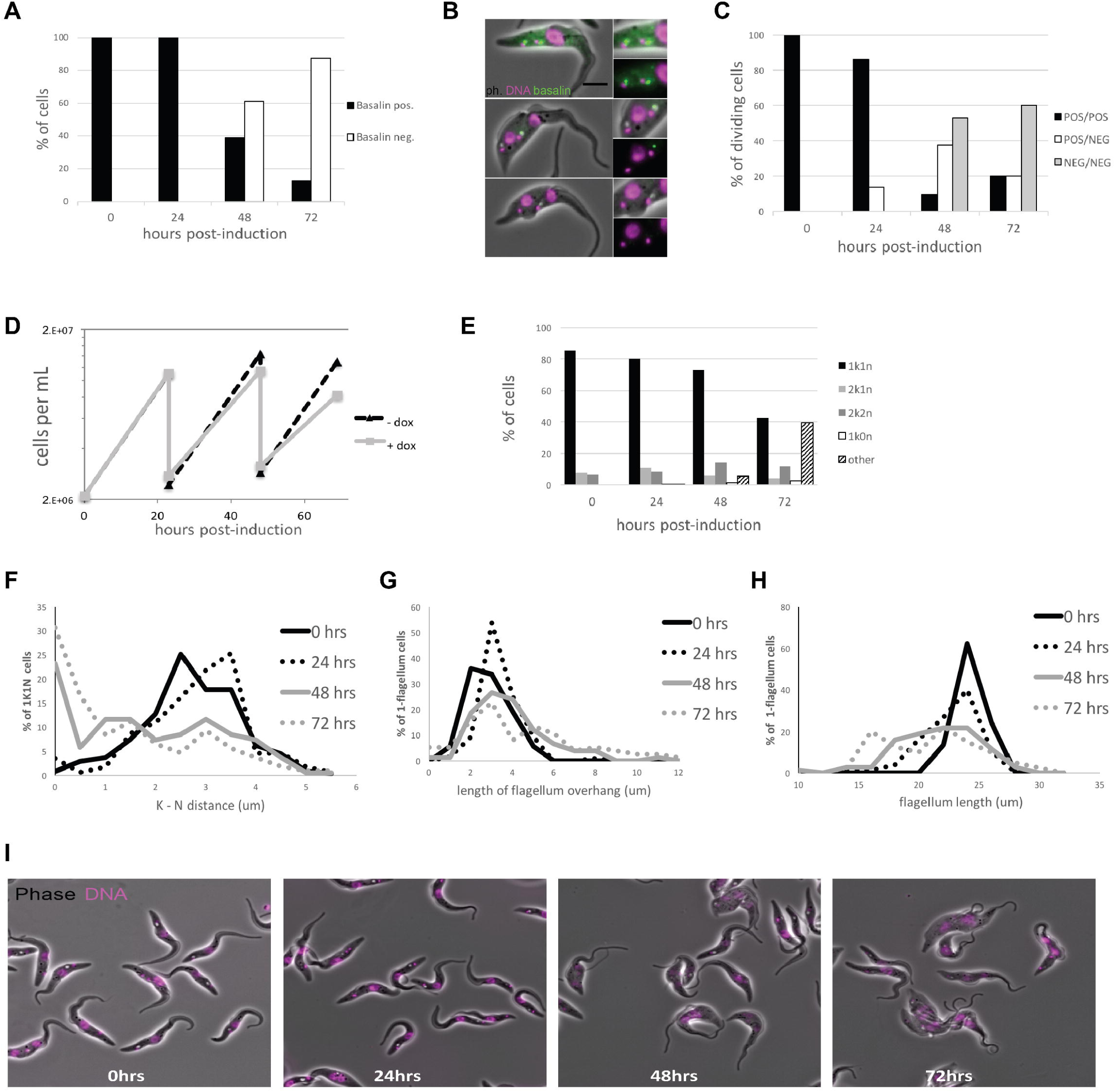
Quantitation of basalin knockdown and phenotype in trypanosome cells. A) Basalin was ablated in cells expressing mNeonGreen∷basalin and cytoskeletons were scored on whether they were positive or negative for basalin at the TZ. RNAi caused ablation such that >80% of cells did not have detectable basalin at 72 hours. B) Examples of dividing cells undergoing basalin RNAi. Non-induced dividing cells have 4 basalin spots that corresponds to localisation at the TZ and pro-basal bodies of both old and new flagella. In cells induced for basalin RNAi where the signal is absent from new flagella, there is only 1 spot that is associated with the old transition zone and the pro-basal body signal is absent. C) Quantitation of basalin signal at the old and new flagella in dividing cells shows that dividing cells with positive old / negative new (POS/NEG) are first observed after 24 hours RNAi and increase in frequency to 60% by 72 hours. D) Growth rates of cells non-induced and induced for basalin RNAi demonstrate that growth rate defect is first observed between 24-48 hours, co-incident with the first cells that are negative for basalin. E) Staining the DNA of cells using Hoechst shows that only the expected kinetoplast-nucleus numbers are observed at 0 hours (i.e. 1k1n, 2k1n, 2k2n), but there is an increase in the proportion of aberrant cells after 48 and 72 hours basalin RNAi. F) There is an accumulation of aberrant morphologies after RNAi. This includes G) a reduction in the kinetoplast-nucleus (k-n) distance, H) an increase in the length of the flagellum that extends beyond the anterior of the cell body (flagellum overhang), and I) a reduction in the overall flagellum length.

**Supplementary Figure 2.**
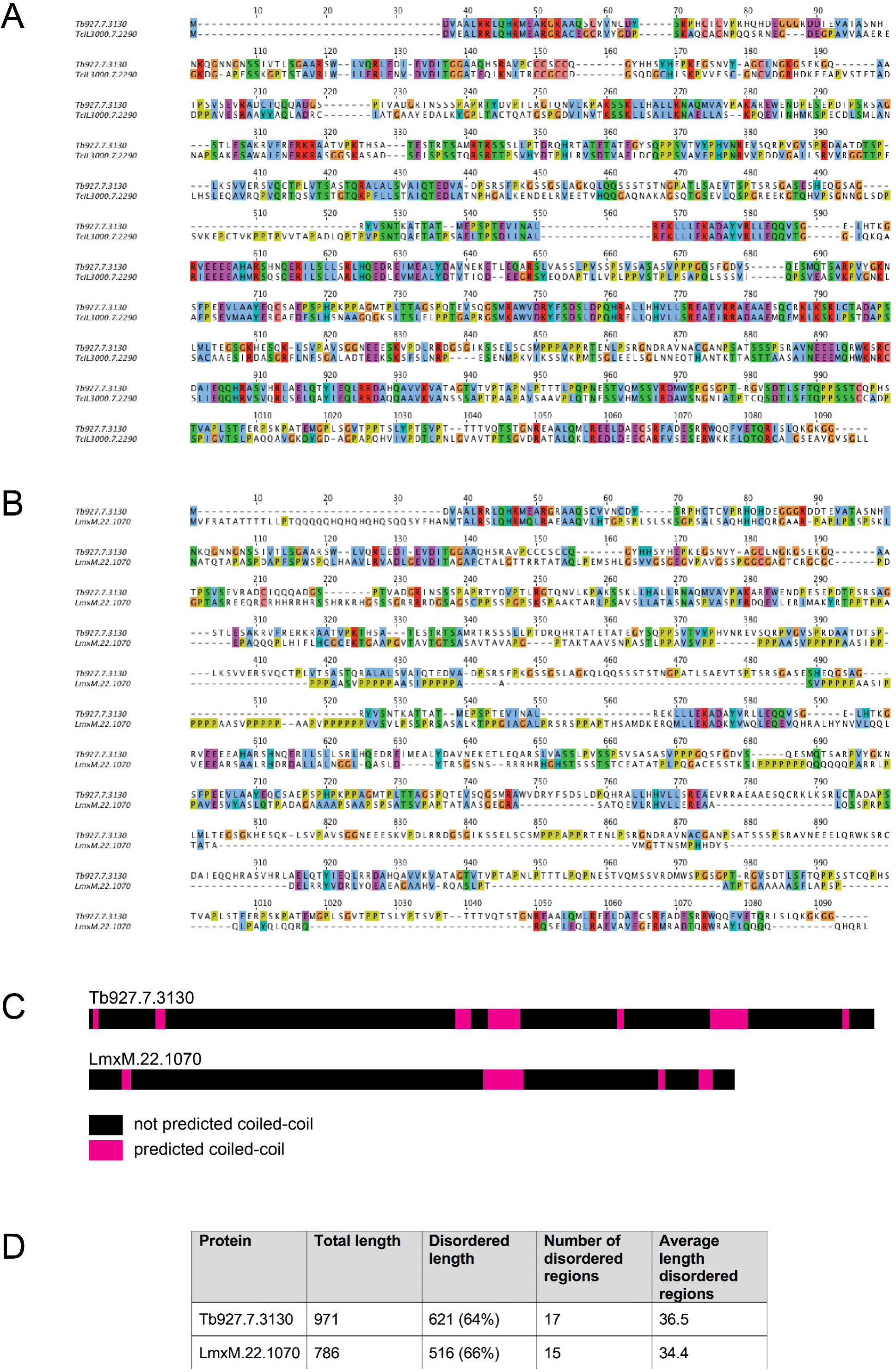
A bioinformatics comparison of Tb927.7.3130 (basalin) and LmxM.22.1070. A) A protein sequence alignment between basalin and its *T. congolense* ortholog. B) A HHpred-driven comparison between Tb927.7.3130 and LmxM.22.1070 demonstrates that most sequence conservation is in the helical segments. C) Both proteins have coiled-coils towards the C terminus of the protein. D) LmxM.22.1070 and basalin show similar levels of intrinsic disorder, with ˜65% disordered residues and 15-17 regions of disorder.

**Supplementary Figure 3.**
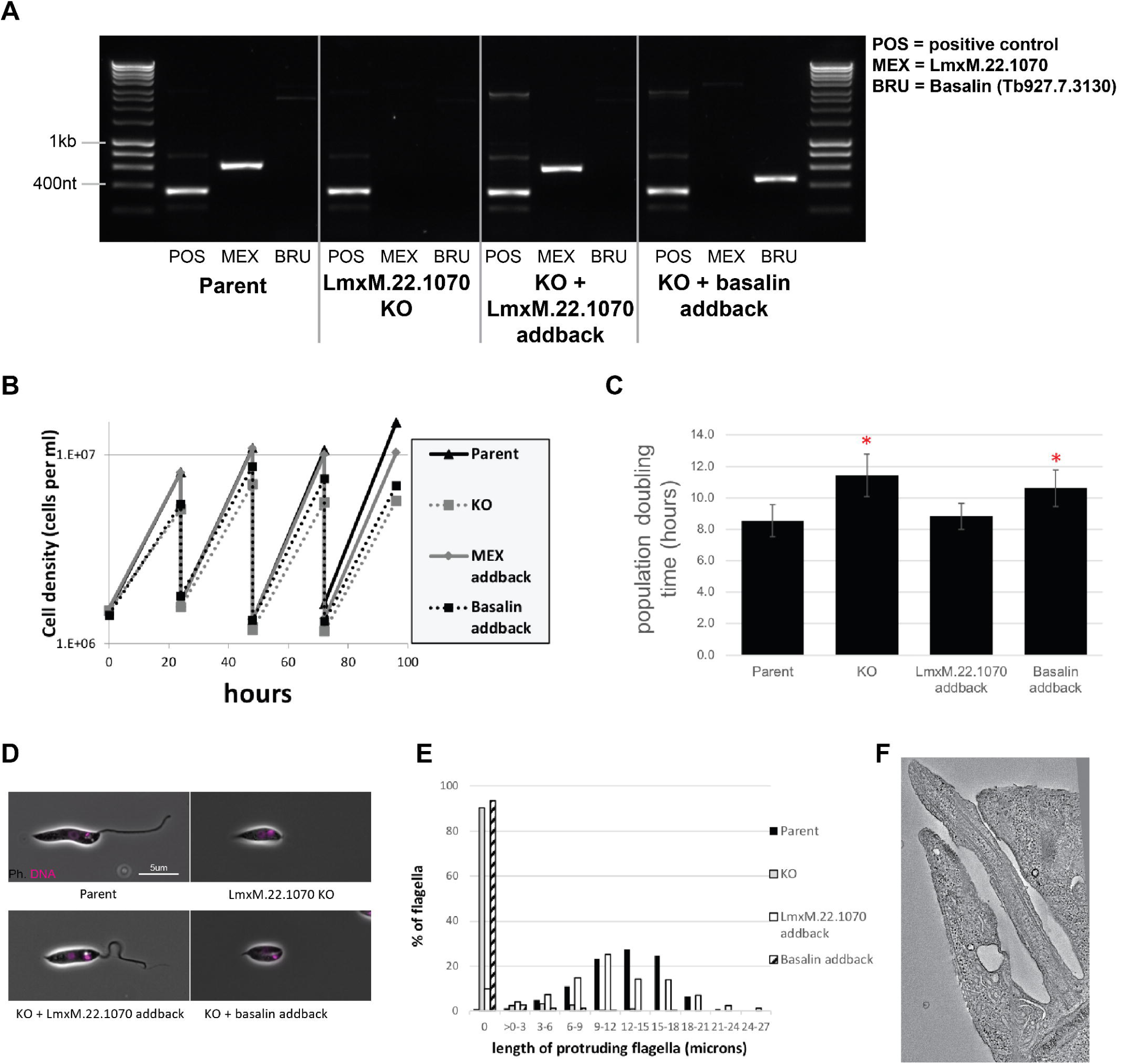
Phenotypic analysis of *Leishmania mexicana* LmxM.22.1070 knockout cells. A) PCR confirms the loss of LmxM.22.1070 in a knockout clone and subsequent addback of LmxM.22.1070 and basalin. B) LmxM.22.1070 knockout cells grow more slowly than the parental cells. Transfection of an episome addback vector containing LmxM.22.1070 rescues this phenotype, but basalin does not. C) The doubling times in of the populations were calculated. The asterisk indicates a statistically significant difference compared to the parental cells (T test, p=0.005). D) Parent and LmxM.22.1070 addback cells have the wildtype morphology, having an elongated cell body and a long flagellum. In contrast, knockout cells and basalin addback cells have a short, round cell body and most cells do not have a flagellum that is visible outside the flagellar pocket of the cell. E) Cytoskeletons prepared from the four cell lines were stained with the TAT anti tubulin antibody to measure the length of the portion of the flagellum that protrudes beyond the flagellar pocket. ˜90% of knockout and basalin addback cells did not have a flagellum that extended beyond the flagellar pocket, and those flagella that did tended to be shorter than the wild type cells. E) A virtual slice through a *Leishmania* LmxM.22.1070 KO tomogram shows that no central pair is evident in the flagellum.

## Bibliography

Aslett, M., Aurrecoechea, C., Berriman, M., Brestelli, J., Brunk, B. P., Carrington, M., et al. (2010). TriTrypDB: a functional genomic resource for the Trypanosomatidae. Nucleic Acids Research, 38(Database issue), D457–62. http://doi.org/10.1093/nar/gkp851

Aurrecoechea, C., Barreto, A., Basenko, E. Y., Brestelli, J., Brunk, B. P., Cade, S., et al. (2017). EuPathDB: the eukaryotic pathogen genomics database resource. Nucleic Acids Research, 45(D1), D581–D591. http://doi.org/10.1093/nar/gkw1105

Beneke, T., Madden, R., Makin, L., Valli, J., Sunter, J., & Gluenz, E. (2017). A CRISPR Cas9 high-throughput genome editing toolkit for kinetoplastids. Royal Society Open Science, 4(5), 170095. http://doi.org/10.1098/rsos.170095

Bindels, D. S., Haarbosch, L., van Weeren, L., Postma, M., Wiese, K. E., Mastop, M., et al. (2017). mScarlet: a bright monomeric red fluorescent protein for cellular imaging. Nature Methods, 14(1), 53–56. http://doi.org/10.1038/nmeth.4074

Branche, C., Kohl, L., Toutirais, G., Buisson, J., Cosson, J., & Bastin, P. (2006). Conserved and specific functions of axoneme components in trypanosome motility. Journal of Cell Science, 119(Pt 16), 3443–3455. http://doi.org/10.1242/jcs.03078

Brun, R., & Schönenberger. (1979). Cultivation and in vitro cloning or procyclic culture forms of Trypanosoma brucei in a semi-defined medium. Short communication. Acta Tropica, 36(3), 289–292. http://doi.org/10.5169/seals-312533

Buday, L., & Tompa, P. (2010). Functional classification of scaffold proteins and related molecules. FEBS Journal, 277(21), 4348–4355. http://doi.org/10.1111/j.1742-4658.2010.07864.x

Callen, D. F., Baker, E. G., & Lane, S. A. (1990). Re-evaluation of GM2346 from a del(16)(q22) to t(4;16)(q35;q22.1). Clinical Genetics, 38(6), 466–468. http://doi.org/10.1111/j.1399-0004.1990.tb03614.x

Collingridge, P. W., & Kelly, S. (2012). MergeAlign: improving multiple sequence alignment performance by dynamic reconstruction of consensus multiple sequence alignments. BMC Bioinformatics, 13(1), 117. http://doi.org/10.1186/1471-2105-13-117

Cumberworth, A., Lamour, G., Babu, M. M., & Gsponer, J. (2013). Promiscuity as a functional trait: intrinsically disordered regions as central players of interactomes. The Biochemical Journal, 454(3), 361–369. http://doi.org/10.1042/BJ20130545

Davey, N. E., Van Roey, K., Weatheritt, R. J., Toedt, G., Uyar, B., Altenberg, B., et al. (2012). Attributes of short linear motifs. Molecular bioSystems, 8(1), 268–281. http://doi.org/10.1039/c1mb05231d

Davy, B. E., & Robinson, M. L. (2003). Congenital hydrocephalus in hy3 mice is caused by a frameshift mutation in Hydin, a large novel gene. Human Molecular Genetics, 12(10), 1163–1170. http://doi.org/10.1093/hmg/ddg122

Dawe, H. R., Shaw, M. K., Farr, H., & Gull, K. (2007). The hydrocephalus inducing gene product, Hydin, positions axonemal central pair microtubules. BMC Biology, 5(1), 33. http://doi.org/10.1186/1741-7007-5-33

Dean, S., Moreira-Leite, F., Varga, V., & Gull, K. (2016). Cilium transition zone proteome reveals compartmentalization and differential dynamics of ciliopathy complexes. Proceedings of the National Academy of Sciences of the United States of America, 113(35), E5135–43. http://doi.org/10.1073/pnas.1604258113

Dean, S., Sunter, J. D., Wheeler, R. J., Hodkinson, I., Gluenz, E., & Gull, K. (2015). A toolkit enabling efficient, scalable and reproducible gene tagging in trypanosomatids. Open Biology, 5(1), 140197. http://doi.org/10.1098/rsob.140197

Dosztányi, Z., Chen, J., Dunker, A. K., Simon, I., & Tompa, P. (2006). Disorder and sequence repeats in hub proteins and their implications for network evolution. Journal of Proteome Research, 5(11), 2985–2995. http://doi.org/10.1021/pr060171o

Dunker, A. K., Cortese, M. S., Romero, P., Iakoucheva, L. M., & Uversky, V. N. (2005). Flexible nets. The roles of intrinsic disorder in protein interaction networks. FEBS Journal, 272(20), 5129–5148. http://doi.org/10.1111/j.1742-4658.2005.04948.x

Dymek, E. E., & Smith, E. F. (2012). PF19 encodes the p60 catalytic subunit of katanin and is required for assembly of the flagellar central apparatus in Chlamydomonas. Journal of Cell Science, 125(14), 3357–3366. http://doi.org/10.1242/jcs.096941

Dymek, E. E., Lefebvre, P. A., & Smith, E. F. (2004). PF15p is the chlamydomonas homologue of the Katanin p80 subunit and is required for assembly of flagellar central microtubules. Eukaryotic Cell, 3(4), 870–879. http://doi.org/10.1128/EC.3.4.870-879.2004

Emms, D. M., & Kelly, S. (2015). OrthoFinder: solving fundamental biases in whole genome comparisons dramatically improves orthogroup inference accuracy. Genome Biology, 16(1), 157. http://doi.org/10.1186/s13059-015-0721-2

Ghedin, E., Bringaud, F., Peterson, J., Myler, P., Berriman, M., Ivens, A., et al. (2004). Gene synteny and evolution of genome architecture in trypanosomatids. Molecular & Biochemical Parasitology, 134(2), 183–191. http://doi.org/10.1016/j.molbiopara.2003.11.012

Guharoy, M., Szabo, B., Contreras Martos, S., Kosol, S., & Tompa, P. (2013). Intrinsic structural disorder in cytoskeletal proteins. Cytoskeleton (Hoboken, NJ), 70(10), 550–571. http://doi.org/10.1002/cm.21118

Höög, J. L., Lacomble, S., O’Toole, E. T., Hoenger, A., McIntosh, J. R., & Gull, K. (2014). Modes of flagellar assembly in Chlamydomonas reinhardtii and Trypanosoma brucei. eLife, 3, e01479. http://doi.org/10.7554/eLife.01479

Inoue, M., Nakamura, Y., Yasuda, K., Yasaka, N., Hara, T., Schnaufer, A., et al. (2005). The 14-3-3 proteins of Trypanosoma brucei function in motility, cytokinesis, and cell cycle. The Journal of Biological Chemistry, 280(14), 14085–14096. http://doi.org/10.1074/jbc.M412336200

Ishida, T., & Kinoshita, K. (2007). PrDOS: prediction of disordered protein regions from amino acid sequence. Nucleic Acids Research, 35(Web Server issue), W460–4. http://doi.org/10.1093/nar/gkm363

Lechtreck, K.-F., & Witman, G. B. (2007). Chlamydomonas reinhardtii hydin is a central pair protein required for flagellar motility. J Cell Biol, 176(4), 473–482. http://doi.org/10.1083/jcb.200611115

Lechtreck, K.-F., Delmotte, P., Robinson, M. L., Sanderson, M. J., & Witman, G. B. (2008). Mutations in Hydin impair ciliary motility in mice. J Cell Biol, 180(3), 633–643. http://doi.org/10.1083/jcb.200710162

Lechtreck, K.-F., Gould, T. J., & Witman, G. B. (2013). Flagellar central pair assembly in Chlamydomonas reinhardtii. Cilia, 2(1), 15. http://doi.org/10.1186/2046-2530-2-15

May-Simera, H. L., & Kelley, M. W. (2012). Cilia, Wnt signaling, and the cytoskeleton. Cilia, 1(1), 7. http://doi.org/10.1186/2046-2530-1-7

McDonnell, A. V., Jiang, T., Keating, A. E., & Berger, B. (2006). Paircoil2: improved prediction of coiled coils from sequence. Bioinformatics (Oxford, England), 22(3), 356–358. http://doi.org/10.1093/bioinformatics/bti797

Mckean, P. G., Baines, A., Vaughan, S., & Gull, K. (2003). Gamma-tubulin functions in the nucleation of a discrete subset of microtubules in the eukaryotic flagellum. Current Biology : CB, 13(7), 598–602. http://doi.org/10.1016/S0960-9822(03)00174-X

Mitchell, D. R., & Sale, W. S. (1999). Characterization of a Chlamydomonas Insertional Mutant that Disrupts Flagellar Central Pair Microtubule-associated Structures. The Journal of Cell Biology, 144(2), 293–304. http://doi.org/10.1083/jcb.144.2.293

Olbrich, H., Schmidts, M., Werner, C., Onoufriadis, A., Loges, N. T., Raidt, J., et al. (2012). Recessive HYDIN mutations cause primary ciliary dyskinesia without randomization of left-right body asymmetry. American Journal of Human Genetics, 91(4), 672–684. http://doi.org/10.1016/j.ajhg.2012.08.016

Overath, P., Haag, J., Lischke, A., & O’hUigin, C. (2001). The surface structure of trypanosomes in relation to their molecular phylogeny. International Journal for Parasitology, 31(5-6), 468–471. http://doi.org/10.1016/S0020-7519(01)00152-7

Poon, S. K., Peacock, L., Gibson, W., Gull, K., & Kelly, S. (2012). A modular and optimized single marker system for generating Trypanosoma brucei cell lines expressing T7 RNA polymerase and the tetracycline repressor. Open Biology, 2(2), 110037. http://doi.org/10.1098/rsob.110037

Redmond, S., Vadivelu, J., & Field, M. C. (2003). RNAit: an automated web-based tool for the selection of RNAi targets in Trypanosoma brucei. Molecular & Biochemical Parasitology, 128(1), 115–118. http://doi.org/10.1016/s0166-6851(03)00045-8

Scott, V., Sherwin, T., & Gull, K. (1997). gamma-tubulin in trypanosomes: molecular characterisation and localisation to multiple and diverse microtubule organising centres. Journal of Cell Science, 110 (Pt 2), 157–168.

Shaner, N. C., Lambert, G. G., Chammas, A., Ni, Y., Cranfill, P. J., Baird, M. A., et al. (2013). A bright monomeric green fluorescent protein derived from Branchiostoma lanceolatum. Nature Methods, 10(5), 407–409. http://doi.org/10.1038/nmeth.2413

Silflow, C. D., Liu, B., LaVoie, M., Richardson, E. A., & Palevitz, B. A. (1999). Gamma-tubulin in Chlamydomonas: characterization of the gene and localization of the gene product in cells. Cell Motility and the Cytoskeleton, 42(4), 285–297. http://doi.org/10.1002/(SICI)1097-0169(1999)42:4<285::AID-CM3>3.0.CO;2-Z

Söding, J. (2005). Protein homology detection by HMM-HMM comparison. Bioinformatics (Oxford, England), 21(7), 951–960. http://doi.org/10.1093/bioinformatics/bti125

Stevens, J. R., & Gibson, W. (1999). The molecular evolution of trypanosomes. Parasitology Today (Personal Ed), 15(11), 432–437. http://doi.org/10.1016/S0169-4758(99)01532-X

Stevens, J., & Rambaut, A. (2001). Evolutionary rate differences in trypanosomes. Infection, Genetics and Evolution : Journal of Molecular Epidemiology and Evolutionary Genetics in Infectious Diseases, 1(2), 143–150. http://doi.org/10.1016/S1567-1348(01)00018-1

Sunter, J. D., & Gull, K. (2017). Shape, form, function and Leishmania pathogenicity: from textbook descriptions to biological understanding. Open Biology, 7(9), 170165. http://doi.org/10.1098/rsob.170165

Uversky, V. N. (2015). The multifaceted roles of intrinsic disorder in protein complexes. FEBS Letters, 589(19 Pt A), 2498–2506. http://doi.org/10.1016/j.febslet.2015.06.004

Zhou, Q., & Li, Z. (2015). γ-Tubulin complex in Trypanosoma brucei: molecular composition, subunit interdependence and requirement for axonemal central pair protein assembly. Molecular Microbiology, 98(4), 667–680. http://doi.org/10.1111/mmi.13149

Zimmermann, L., Stephens, A., Nam, S.-Z., Rau, D., Kübler, J., Lozajic, M., et al. (2018). A Completely Reimplemented MPI Bioinformatics Toolkit with a New HHpred Server at its Core. Journal of Molecular Biology, 430(15), 2237–2243. http://doi.org/10.1016/j.jmb.2017.12.007

